# A rare deletion of *IhUF3GT1* in the ivy-leaf morning glory eliminates anthocyanin-based pigmentation

**DOI:** 10.64898/2026.06.08.730875

**Authors:** Damian J. Hernandez, Emily Glasgow, Margaret C. Li, Martin R. Henry, Amanda L. Peake, John R. Stinchcombe

**Author notes:** These authors contributed equally (Damian J. Hernandez, Emily Glasgow). These authors contributed equally (Margaret C. Li, Martin R. Henry).

## Abstract

Anthocyanin biosynthesis is a highly branched network with upstream mutations leading to the loss of many other branches beyond just anthocyanins like flavonols, isoflavonoids, and proanthocyanidins. Consequently, disentangling the specific effects of anthocyanins on plant fitness from those of other branches is difficult unless a mutant impacting the terminal step of anthocyanin biosynthesis can be found. We discovered an *Ipomoea hederacea* plant (ivy-leaf morning glory) lacking anthocyanins due to a genetic deletion in the terminal step of the anthocyanin biosynthetic pathway. Anthocyanin loss follows a recessive, Mendelian inheritance pattern. Anthocyanin loss is perfectly correlated with a lack of expression of the anthocyanin biosynthesis gene *IhUF3GT1 (anthocyanidin-3-O-glucosyltransferase)*. *IhUF3GT1* is the gene most strongly and consistently differentially expressed between unpigmented and pigmented flowers in F_2_ siblings. A deletion of half of the *IhUF3GT1* gene which includes the start codon is the genetic basis underlying a loss of *IhUF3GT1* expression and, consequently, anthocyanins in unpigmented plants. The *IhUF3GT1* deletion is rare and unique to the unpigmented line. Genome coverage analyses demonstrate that no other pigmented *I. hederacea* line (of 123 screened) has a loss-of-function mutation in *IhUF3GT1*. Furthermore, we confirm cosegregation of the *IhUF3GT1* deletion with the unpigmented phenotype in F_3_ progeny from a cross where all unpigmented plants are homozygous for the *IhUF3GT1* deletion. Taken together, we describe the genetic basis of a novel unpigmented mutant in *I. hederacea*, creating a potentially important model for studying the fitness effects of anthocyanin loss.

## INTRODUCTION

Anthocyanins are plant pigments underlying important traits in plants like floral color (Rausher, 2008) and tolerance to high levels of UV radiation (Yan *et al*., 2022). For example, anthocyanin synthesis genes are also often upregulated in plants experiencing drought, enhancing resilience to osmotic stress by scavenging cell-death-inducing H_2_O_2_ (Ma *et al*., 2025). Anthocyanin biosynthesis also serves as an important focal trait for understanding the inheritance and evolution of ecologically important traits (Strauss and Whittall, 2006; Baucom *et al*., 2011; Smith *et al*., 2013). Mutations in different anthocyanin synthesis genes can create similar phenotypes in natural populations because the anthocyanin biosynthesis pathway has multiple potential points of failure (Wu *et al*., 2013; Coburn *et al*., 2015), leading to multiple independent genetic bases for similar phenotypes. Consequently, the anthocyanin pathway has become a model for connecting the genetic mechanisms underlying phenotypic variation with their ecological and evolutionary consequences, replicated across species or independent mutations in the same species (Rausher *et al*., 1993; Clegg and Durbin, 2000; Fehr and Rausher, 2004; Twyford *et al*., 2018). Here, we characterized a phenotypically and genetically rare variant of anthocyanin-based pigmentation in *Ipomoea hederacea* (ivy-leaved morning glory) which leads to a dramatic shift from vibrant, blue/purple flowers and pigmented stems to a complete loss of blue/purple pigmentation throughout the whole plant (Figure 1).

**Figure 1.**
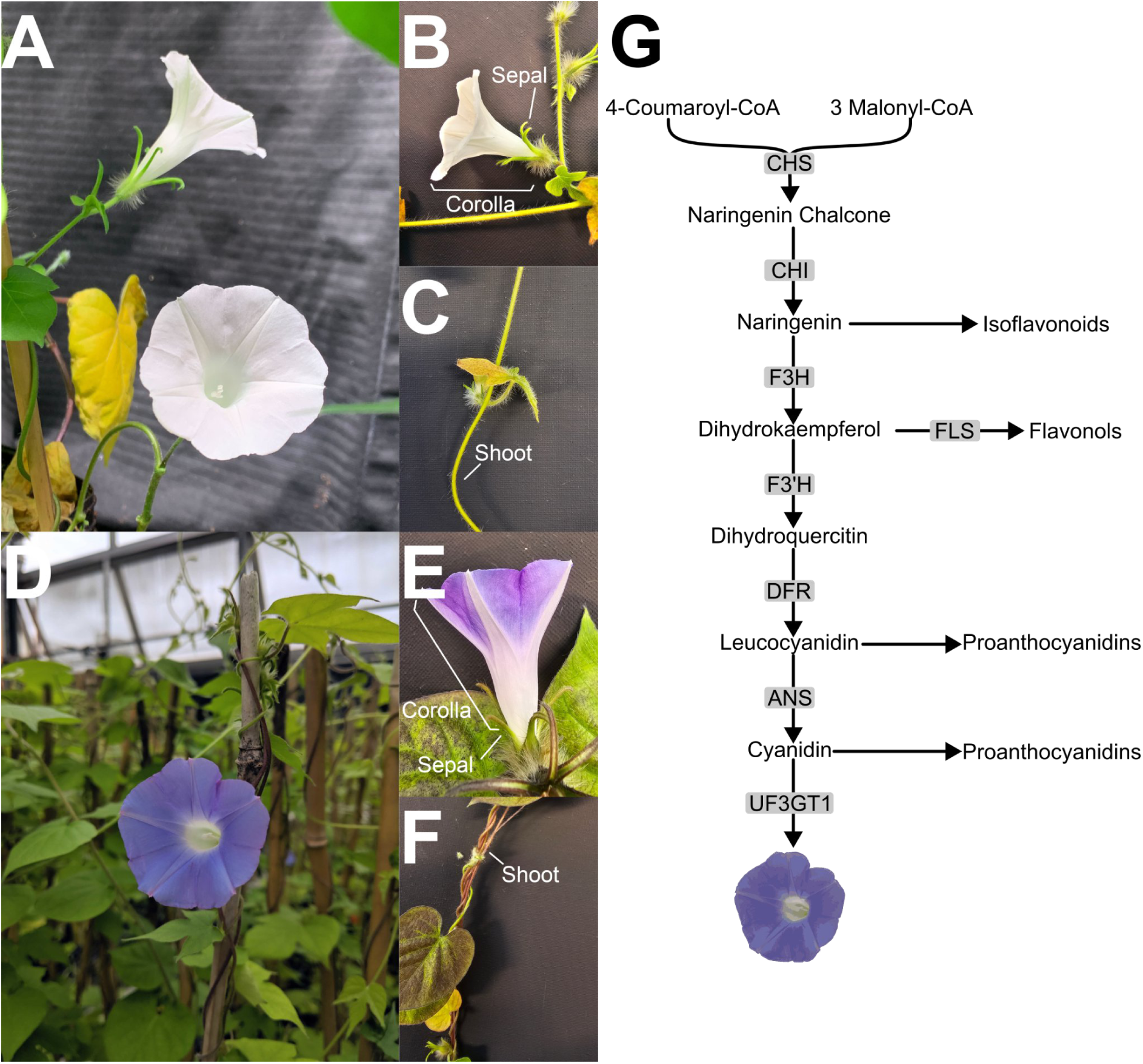
Lack of pigmentation is a complete loss throughout the whole plant. A) Panned-out view of mutant plants lacking anthocyanin pigments. Flowers are completely white and the vegetative portions of the plant are green indicating a lack of anthocyanin-derived pigments. B) Side-view of flowers demonstrating lack of anthocyanin-derived pigmentation in either the petals or sepals. C) Magnified view of morning glory shoot showing a lack of anthocyanin-derived pigments in the vegetative portions of the plant. D) Panned-out view of canonical *I. hederacea* plants with blue/purple flowers and pigmented shoots. E) Side-view of canonical *I. hederacea* flowers showing anthocyanin-derived pigments in both petals and sepals. F) Magnified-view of canonical *I. hederacea* shoots demonstrating intense pigmentation of shoots. G) Cartoon schematic of how loss of *IhUF3GT1* leads to loss of pigmentation. The only anthocyanin branch to produce pigments in *I. hederacea* is expected to be the cyanidin branch (blue/purple pigments). The delphinidin branch of anthocyanin production is absent in *Ipomoea* (Rausher, 2006). Red/orange-pigments (pelargonidin branch) have evolved at least four times in *Ipomoea*, but not in the lineage in which *I. hederacea* is present (Streisfeld and Rausher, 2009). CHS: Chalcone synthase, CHI: Chalcone isomerase, FLS: Flavonol synthase, F3H: Naringenin 3-dioxygenase, F3’H: Flavonoid 3’,5’-hydroxylase, DFR: Dihydroflavonol 4-reductase/flavanone 4-reductase, ANS: Anthocyanidin synthase, *IhUF3GT1*: Anthocyanidin 3-O-glucosyltransferase.

Mutations at several points in the anthocyanin pathway create similar phenotypes where plants/species often transition from predominantly blue/purple pigments to red or no pigments (Rausher, 2008; Ho and Smith, 2016; Marin-Recinos and Pucker, 2024). For example, loss or down-regulation of either *CHS*, *CHI*, or *DFR* eliminates biosynthesis of anthocyanins, with each loss-of-function mutation or down-regulation producing white flowers (e.g., in the common morning glory *I. purpurea*; Fukada-Tanaka *et al*., 1997; Johzuka-Hisatomi *et al*., 1999; Coberly and Rausher, 2003). However, anthocyanins are only one branch of a more complex network of flavonoid synthesis like the generation of proanthocyanidins (Dixon *et al*., 2005). Consequently, loss-of-function mutations that disrupt anthocyanin synthesis at an early stage in the pathway like *CHS* may impact other traits (Figure 1G) like herbivory defense through proanthocyanidins (Forkner *et al*., 2004). As a result, mutations at different parts of the pathway are likely to have different pleiotropic consequences, and evolutionary dynamics.

Intensive investigation of the molecular and phenotypic evolution of anthocyanin variation, especially with regard to flower color, has yielded mixed results about the evolutionary rate of different portions of the pathway. Early molecular population genetics studies suggested that regulatory loci evolve faster than the structural genes of the pathway (Tiffin *et al*., 1998; Rausher *et al*., 1999) and that upstream genes in the pathway evolved more slowly than downstream genes (Rausher *et al*., 1999). Subsequent work, however, with more species found no support for reduced evolutionary rates in upstream enzymes in the pathway (Wheeler *et al*., 2022). However, Streisfeld and Rausher (2011) found that mutations involving transcription factors and cis-regulatory mutations for anthocyanin pathway genes had higher fixation rates than coding mutations in anthocyanin pathway genes, presumably due to the pleiotropic costs of the latter.

At the phenotypic level, numerous species have intermediate frequency anthocyanin-based flower color polymorphisms (Warren and Mackenzie, 2001; Del Valle *et al*., 2019), show phenotypic clines (Dick *et al*., 2011; Maunder *et al*., 2026), and show some evidence of being actively maintained by selection (Subramaniam and Rausher, 2000). Phenotypic studies suggest greater fitness consequences of mutations to structural rather than regulatory genes in the pathway. For example, the *a* allele in *I. purpurea* produces white flowers when homozygous (*aa*), due to a transposable element insertion in *CHS-D* (Habu *et al., 1998*), appears to be at low frequency in natural populations (<0.01; Fehr and Rausher, 2004; Coberly and Rausher, 2008), associated with lower survival to flowering in the field (Coberly and Rausher, 2008), and lower fecundity at high temperatures (Coberly and Rausher, 2003). In contrast, multiple investigations of the *w* allele, which also leads to white flowers in the *ww* genotype in *I. purpurea* (due to inactivation of the transcription factor *IpMyb1* (Chang *et al*., 2005) have failed to reveal strong fitness costs (Rausher *et al*., 1993; Rausher and Fry, 1993; Fineblum and Rausher, 1997; Fry and Rausher, 1997)).

Whole-plant loss of anthocyanins and loss-of-function mutations in enzymes of the pathway (Figure 1), often appear to be rare in populations. For example, the frequency of ‘albino’ individuals of *Delphinium nelsonii* appears to be <0.1%, and they show reduced pollinator visitation and seed set compared to blue–flowered individuals, leading Waser and Price (1981) to interpret their frequency as the result of mutation-selection balance. Whole-plant anthocyanin loss occurs at very low frequency in *Silene littorea* (<1%; Del Valle *et al*., 2019), consistent with several other species they reviewed. Reports from *Mimulus lewisii* and *Iochroma calycinum* both implicate mutations to *DFR* in producing white flowers and anthocyanin loss (Wu *et al*., 2013; Coburn *et al*., 2015), and both are reported to be rare in populations. One exception to this general pattern is from *Mimulus guttatus*, where complete loss of anthocyanin is at high frequency (>20%); mutants showed few fitness costs, and had later flowering time and greater investment in asexual reproduction, presumably due to pleiotropy or linkage (Twyford *et al*., 2018).

Collectively, these studies suggest mixed results on the rates of molecular evolution of anthocyanin pathway genes, with some hints of pleiotropy leading to differential fixation rates; differences in allele frequencies and demonstrated pleiotropic, fitness costs of mutations producing phenotypically similar white flowers; and an overall rarity of both coding sequence mutations in the enzymatic genes of the pathway and whole-plant loss of anthocyanins. The diverse potential functions of anthocyanins across plant lineages, and thus consequences of their loss, may be one reason for these mixed results from population and ecological genetics.

To date, the ivy-leaf morning glory, *I. hederacea* has been exclusively reported to have blue/purple pigmentation (i.e., cyanidin-derived pigments). However, we found one maternal family of *I. hederacea* previously collected by Henry and Stinchcombe (2023a) with white flowers that also lack any visible anthocyanin-derived pigments throughout the whole plant (Figure 1). We combined test crosses, transcriptomics, genome-coverage, and co-segregation analyses to: 1) demonstrate that the unpigmented phenotype is a Mendelian trait with a homozygous and recessive inheritance pattern, 2) establish that deletion of the anthocyanin-producing gene *IhUF3GT1* is the genetic basis underlying the unpigmented phenotype, and 3) loss of anthocyanins stems from a single structural variant in which half of *IhUF3GT1* is missing. Taken together, we discover a previously undescribed mutation in the *Ipomoea* genus that completely eliminates biosynthesis of anthocyanins.

## METHODS

### Study system

The ivy-leaf morning glory, *Ipomoea hederacea* (Convolvulaceae) is an annual vine commonly found in disturbed habitats in the southeastern and mid-Atlantic U.S.A. Plants germinate in early summer, often following disturbance, and grow and flower until a killing frost. Despite showy blue/purple flowers, selfing rates are high (93%, Ennos, 1981; 92-94%, Campitelli, and Stinchcombe, 2014).

### Plant-crosses: generating hybrids between white- and purple-flowered parents

Plants were grown at the University of Toronto St. George Greenhouse in conical containers with Sunshine Potting Mix (cat. no.: 5101.CFL002.8). We grew plants at 26-30℃ and 16:8 day:night light cycles for 45 days. Two weeks post-germination, we applied 4 mg/mL of 20-20-20 PlantProd Classics fertilizer (cat. no.: 10529). After this initial fertilization, we provided plants with 4 mg/mL weekly of Miracle Gro Bloom Booster 15-30-15 (cat. no.: 2756210), unless indicated otherwise below. To initiate flowering, we grew plants at 20-24℃ and 8:16 day:night light cycles.

We grew one seed each from a pigmented (MD462; 38.83662°N, -76.07692°W; Easton, Maryland; Campitelli and Stinchcombe, 2013b), and unpigmented line (401S; 35.091722°N, 79.742722°W; Ellerbe, North Carolina; Henry and Stinchcombe, 2023a). Once these maternal lines flowered, we removed the immature anthers of the pigmented plant in an unopened flower bud and transferred pollen from the unpigmented parent to the pigmented parent’s stigma with forceps. To prevent cross-contamination, we cleaned forceps with ethanol. We obtained 13 successful crosses. We dried these F_1_ seeds in the laboratory for one month.

After drying, two seeds per cross were scarified by physically nicking the seed coat with a clean blade to allow water to enter the seed coat, and scarified seeds were planted individually in 4 L pots containing Sunshine potting mix in a growth room. Pots were watered as needed, and fertilizing was carried out as above. Plants were allowed to self-pollinate, and F_2_ seeds were collected as above. Progeny from the F_1_ individual that set the largest number of seeds was used for the next phase of the experiment (i.e., all progeny are derived from a single F_1_ cross).

After drying, 500 F_2_ seeds were planted in 1L rectangular pots (one seed per pot). Two weeks post-germination, 20-20-20 PlantProd Classics fertilizer was applied at a rate of 4 mg/mL; subsequently, slow-release phosphorus and potassium were applied to provide fertilizer for the remainder of the experiment. Of the 500 initially planted, 494 survived for quantification and seed collection (to produce F_3_ seed).

### RNA-Seq and differential expression between white and purple flowers

From the F_2_ population described above, we extracted RNA from 100 mg of petals from four white-flowered plants and four purple-flowered plants. Flowers were collected in the evening at the same time before opening. RNA was extracted from flowers with the Spectrum Plant Total RNA Kit (Sigma-Aldrich, STRN50). Paired-end libraries were prepared using the NEBNext Ultra II Directional RNA-Poly(A) Library Prep for Illumina (New England Biolabs, E7760) with 300 ng of input RNA. Libraries were sequenced on a 10B NovaSeq X flow cell (150 bp reads) at The Centre for Applied Genomics in The Hospital for Sick Children (Toronto, ON, Canada).

We decompressed RNA-Seq data with Illumina’s DRAGEN ORA software (v2.7.0, Illumina). We assessed the sequencing quality of the raw decompressed reads using *FastQC* (v0.12.1; Andrews, 2010) and *MultiQC* (v1.30; Ewels *et al*., 2016). We initially conducted soft-trimming of raw reads using *TrimGalore* (0.6.10; Krueger *et al*., 2023) with the default PHRED score cut-off of 20. Based on sequencing quality of the trimmed reads, we chose to clip our reads at the 5’ and 3’ ends by 5 and 10 bp, respectively, after soft-trimming. This clipping did not change our results.

We aligned reads to a recent assembly of *Ipomoea hederacea* (Peake *et al*., 2026) using the *STAR* aligner (v2.7.11b; Dobin *et al*., 2013). To identify differentially-expressed genes, we compared gene expression between white and purple flowers with *DESeq2* (v1.48.1; Love *et al*., 2014) using Likelihood-Ratio Tests in which we compared models with flower color as a predictor to null models without. We corrected for multiple comparisons using a Benjamini-Hochberg correction and used an FDR cut-off of 0.05 to determine significance. To identify gene function, we mapped all *I. hederacea* genes to *Arabidopsis thaliana* (Araport11; Cheng *et al*., 2017) and *Solanum lycopersicum* (ITAG4.0; Hosmani *et al*., 2019) proteomes using *DIAMOND*’s (v2.1.13; Buchfink *et al*., 2015, 2021) *blastp* function and “more-sensitive” mode. We determined if *I. hederacea* genes are involved in anthocyanin production by confirming they match *S. lycopersicum* genes in a curated subset of the flavonoid (ko00941) and anthocyanin (ko00942) biosynthesis pathways in the Kyoto Encyclopedia of Genes and Genomes (KEGG; Kanehisa *et al*., 2016). *3’GT* only has a representative sequence in KEGG for *Gentiana triflora*; thus, we searched the *I. hederacea* genome for the *G. triflora 3’GT* sequence.

### Genome coverage analyses of pigmented and unpigmented plants

We next generated a genome sequence of the unpigmented parent, as a complement to a previously described sample of 123 genome sequences from naturally pigmented plants, including the pigmented parent (Peake *et al*., 2026). We extracted genomic DNA from leaf tissue of the unpigmented parent using the Qiagen DNeasy Plant Mini Kit (Qiagen, 69106), and library preparation and short-read sequencing was performed by The Centre for Applied Genomics in The Hospital for Sick Children (Toronto, ON, Canada). Libraries were prepared with 5200 ng of DNA using the Nextera XT DNA Library Preparation Kit kit (Illumina, FC-131). We performed short-read resequencing on an Illumina Nova Seq X platform using paired-end sequencing (2x150 bp).

We aligned reads to the *I. hederacea* reference genome (Peake *et al*., 2026), removed adapter sequences using *TrimGalore (Krueger et al., 2023)*, mapped reads to the *I. hederacea* genome using *BWA-MEM2* (v2.2.1; Vasimuddin *et al*., 2019), and sorted and indexed reads using *samtools* (v1.22.1; Li *et al*., 2009; Danecek *et al*., 2021). The unpigmented parent was sequenced to a depth of 260,258,530 reads. We next compared sequence coverage of the white parent to 123 naturally pigmented lines collected from *I. hederacea*’s range across the United States of America. We calculated coverage using *mosdepth* (v0.3.3; Pedersen and Quinlan, 2018) within 100 bp windows and filtered for reads with a minimum mapping quality of 30.

### Co-segregation of IhUF3GT1 loss with an unpigmented phenotype

Preliminary analysis suggested a loss of *IhUF3GT1* was associated with a loss of pigmentation (*see below*). To determine if loss of *IhUF3GT1* underpins the loss of pigmentation, we grew one F_3_ seed set by a sample of pigmented and unpigmented F_2_ individuals to validate the association between *IhUF3GT1* and pigmentation. We screened 17 unpigmented and 36 pigmented F_3_ plants for the presence or absence of the *IhUF3GT1* locus. We extracted genomic DNA from plants using the Qiagen DNeasy Plant Mini Kit (Qiagen, 69106). We determined if pigment loss co-segregates with *IhUF3GT1* loss by PCR using the primers listed in Table 1. As controls, we used genomic DNA from full siblings of the unpigmented and pigmented parents from the F_1_ cross above. DNA from the unpigmented sibling was the biological negative, DNA from the pigmented sibling was the biological positive, and sterile water was the reaction negative. As reaction positive controls, we used another set of *I. hederacea* primers (IH00534) from Campitelli, and Stinchcombe, 2014. For the reaction, 1 μL of genomic DNA was used (∼40 ng) in a 25 μL reaction following manufacturer’s guidelines (GoTaq Green Master Mix, Promega, M7123). The PCR was run in a thermocycler with the settings in Supplementary File 1. PCR products were visualized on 1% agarose gels and stained with SafeView Classic (Diamed, ABMG108). This assay is a presence/absence genotyping assay in which a positive signal for *IhUF3GT1* indicates that that plant has the gene and a negative signal indicates it is missing the gene/region.

**Table 1.**
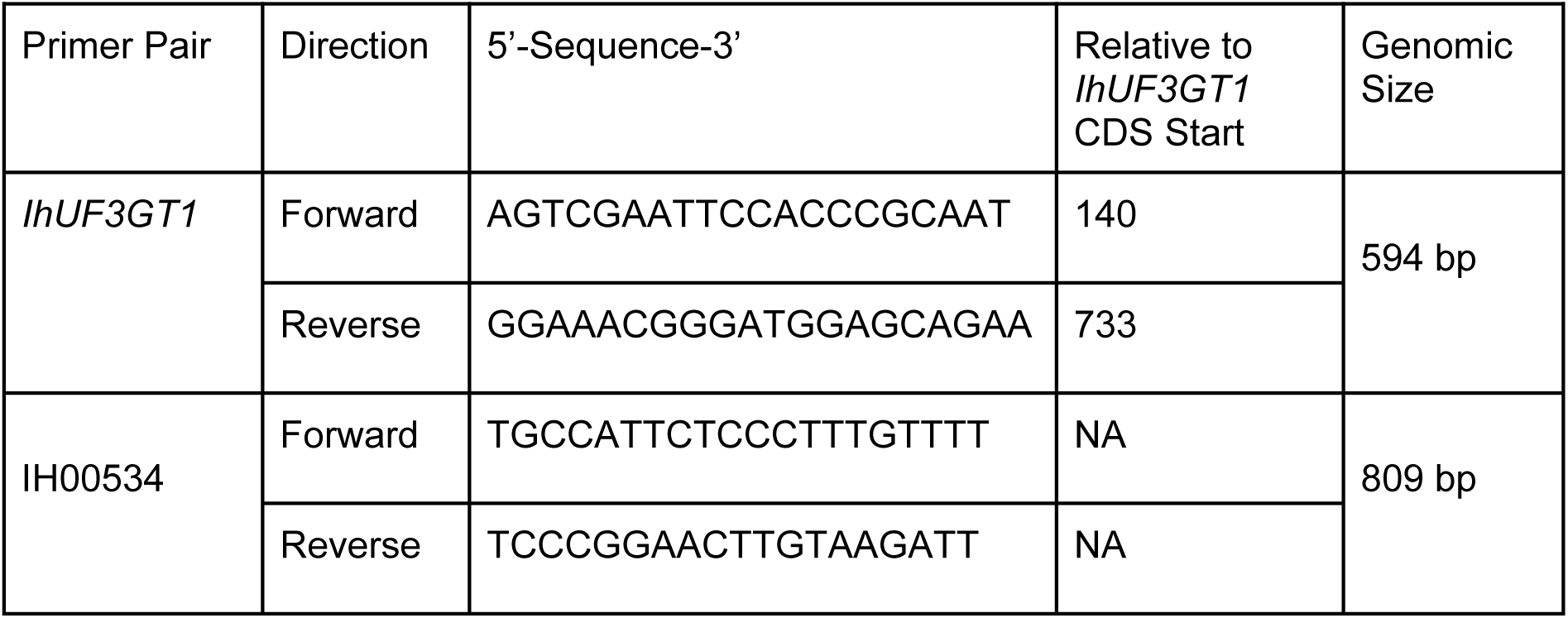
Primers used in co-segregation analysis. “*IhUF3GT1* CDS” Start represents where the 5’ end of the primer binds along *IhUF3GT1*’s coding sequence. The start of the *IhUF3GT1* deletion in the unpigmented parent is position 733 in the *IhUF3GT1* coding sequence. The *IhUF3GT1* primers span the start of the deletion. *IH00534* primers are microsatellite markers from (Campitelli, and Stinchcombe, 2014) which serve as a positive reaction control.

### Data availability

Sequencing data for the white flower parent genome and gene expression are deposited in NCBI (BioProject: PRJNA1355176, SRA: SRP638260). Our code and computational environments will be made available on GitHub following publication, a reviewer link is currently available at: https://zenodo.org/records/20596819?preview=1&token=eyJhbGciOiJIUzUxMiJ9.eyJpZCI6Ij k2NGUxZmRlLWExOGItNDZkMS1iMjhiLTQ0OGI5ODljNmZjMiIsImRhdGEiOnt9LCJyYW5kb 20iOiI4MGQ2NTQzZGExMzliYjZkZDFjNmY2YjIxY2EwZDBjYSJ9.PbK7Dz4dhHcHDKbUni7 WP7PBvcz3WWgastCe3V2etyCeRAPeQuOUS5jGFYMWPsQ2f2l289uXF0bYPEo5h-mBEg. Packages used to conduct analyses in *R* (v4.5.1; R Core Team, 2025) are also listed in Supplementary File 1.

## RESULTS

### Absence of blue pigments is heritable and recessive

We crossed *I. hederacea* plants with and without pigmentation. In the F_2_ population, 28.5% of offspring (141/494 plants, Figure 2) were unpigmented, indicating that the unpigmented phenotype is homozygous, recessive, and likely determined by a single locus (χ^2^-test, 3:1 expectation; χ^2^ = 3.3063, p = 0.07). We observed a slight excess of unpigmented offspring relative to expectations (28.5% versus 25%), but not enough to produce a significant difference from a 1-locus model.

**Figure 2.**
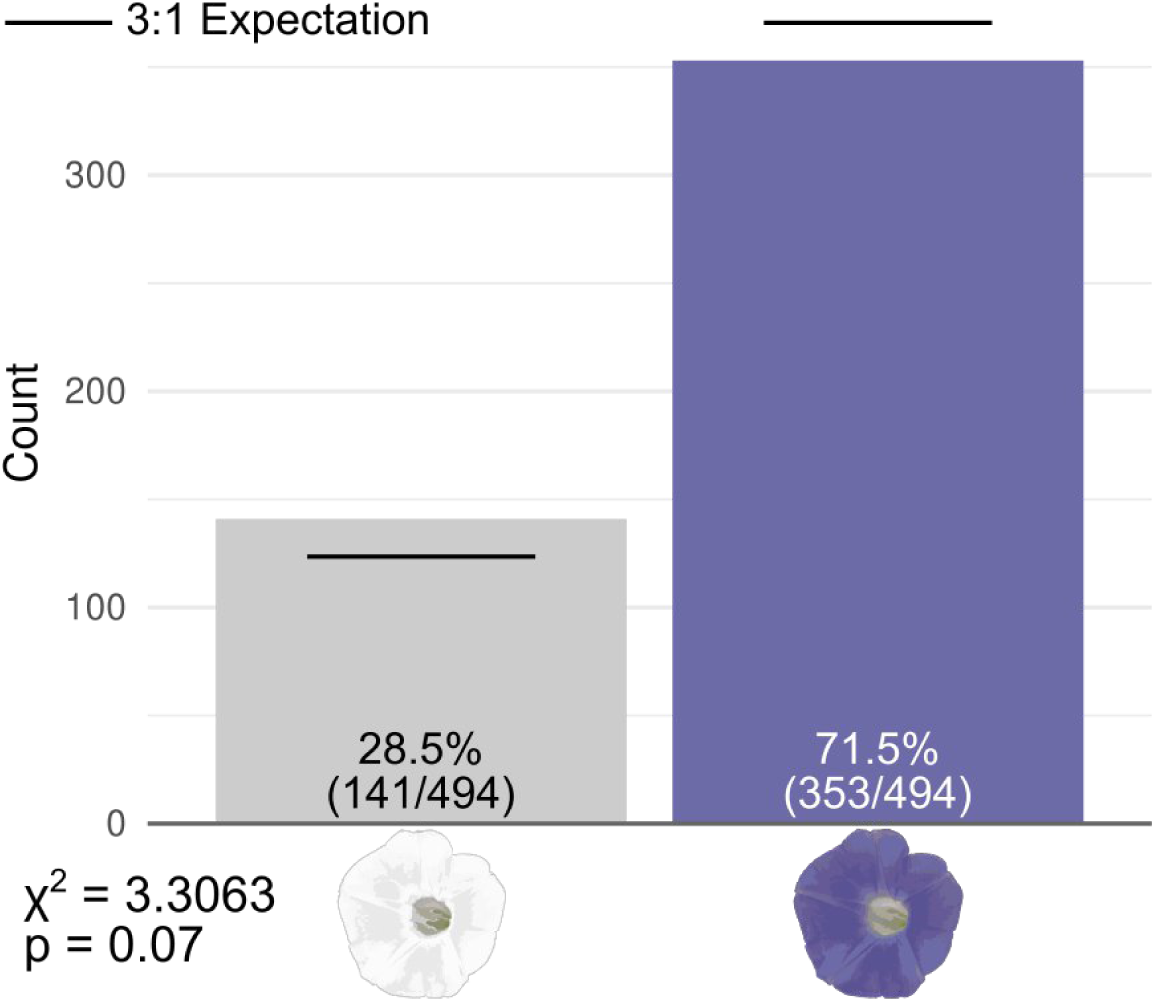
Pigment loss follows a homozygous, recessive inheritance pattern. Of 494 plants from an F_2_ cross between unpigmented and pigmented parents, 28.5% (141) were unpigmented (gray bar) and 71.5% (353) were pigmented (purple bar). These frequencies do not deviate significantly from the expectations of a Mendelian trait (3:1, black lines) in a well powered experiment (N = 494 plants). We did observe slightly more white flowered plants (and fewer pigmented plants) than expected under simple Mendelian inheritance, but this difference was not significant (X^2^ = 3.3063, p = 0.07). Thus our results strongly support that the unpigmented phenotype is a single-locus, homozygous, recessive trait.

Additionally, having white flowers completely co-occurs with a lack of pigmentation in *I. hederacea* stems. A haphazard sample of 44 plants displaying white flowers indicated that all of them were also unpigmented in their stems, strongly suggesting that the genetic basis of the unpigmented phenotype is a complete loss of the ability to synthesize anthocyanin-based pigments rather than a difference in cell-type expression (e.g., as with pigment intensity and pattern in *I. purpurea*; (Park *et al*., 2007)).

### White flowers do not express the essential anthocyanin-producing gene IhUF3GT1

Expression of the *I. hederacea* gene *g2417* is absent in white flowers and present in purple flowers (Figure 3), with absence of *g2417* expression being the single most striking transcriptomic difference between white and purple flowers. *g2417* most closely matches the *anthocyanidin 3-O-glucosyltransferase* gene, *IhUF3GT1*, in tomato (*Solanum lycopersicum,* ITAG4.0, e-value = 1.21 ✕ 10^-151^; common alternative names: *BZ1*, *UF3GT*, *UFGT*, *U3GT*). *IhUF3GT1* is essential in producing pigmented flowers in other plant species by converting anthocyanidins to anthocyanins (Rausher, 2006). No other genes in the anthocyanin-biosynthesis pathway were differentially expressed between white and purple flowers (Figure 3B). Taken together, these results suggest that conversion of anthocyanidins like cyanidin to their anthocyanin derivatives is the main mechanism underlying a lack of pigment in white flowers.

**Figure 3.**
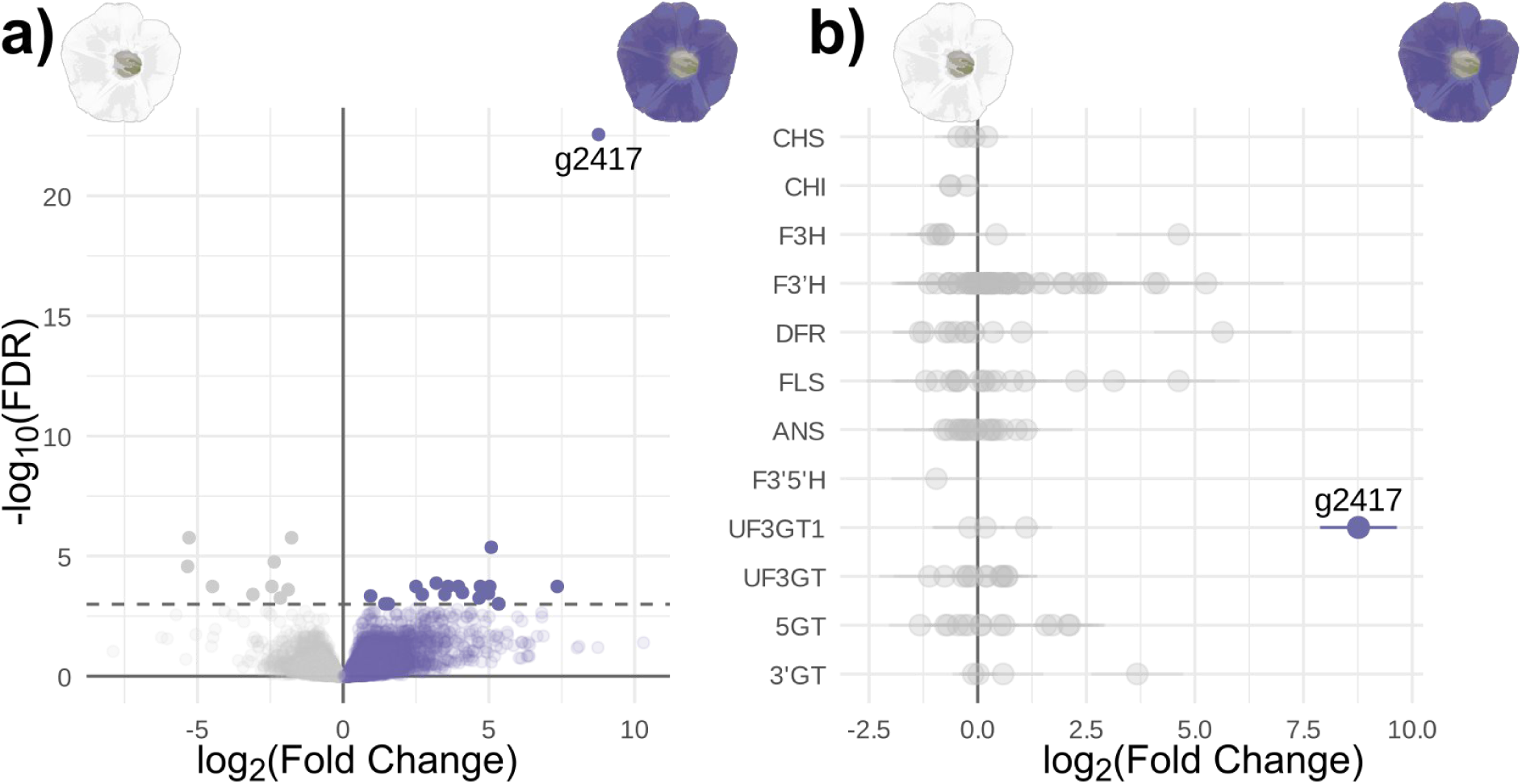
Unpigmented flowers do not express the anthocyanin-producing gene *IhUF3GT1*. a) Differential expression of all expressed genes in white (unpigmented, left) vs purple (pigmented, right) flowers. Solid colors represent significantly differentially expressed genes (FDR < 0.05). Purple points represent genes with higher expression in purple flowers. Grey points represent genes with higher expression in white flowers. The dashed line is the FDR cut-off of 0.05. b) Fold change in expression (white vs purple flowers) of 10 key genes in the anthocyanin biosynthesis pathway present in *I. hederacea*. Genes are sorted by their position in the pathway (higher genes are upstream). Faded grey points are not differentially-expressed (FDR > 0.05). Error bars are the standard error in fold change from negative binomial regressions in *DESeq2*. *g2417* (solid purple) is the only gene in the anthocyanin biosynthesis pathway that is differentially expressed between white and purple flowers.

### A large deletion at the IhUF3GT1 locus underlies loss of pigmentation

Given the association between pigment loss in white flowers and an absence of *IhUF3GT1* expression, we hypothesized that the lack of *IhUF3GT1* expression was due to genetic changes like nonsense mutations or large deletions. We thus conducted a genome coverage analysis in which we assessed the depth of sequencing coverage at the *IhUF3GT1* locus in 123 *I. hederacea* lines.

The parental, unpigmented line has an up to 7 kilobase deletion in the region within and upstream of the *IhUF3GT1* gene (Figure 4A). The *IhUF3GT1* deletion removes 50.3% of the coding sequence, including the start codon (Figure 4A). *IhUF3GT1* is present in all other pigmented lines (n = 123; Figure 4B), indicating that *IhUF3GT1* deletion is unique to the unpigmented phenotype.

**Figure 4.**
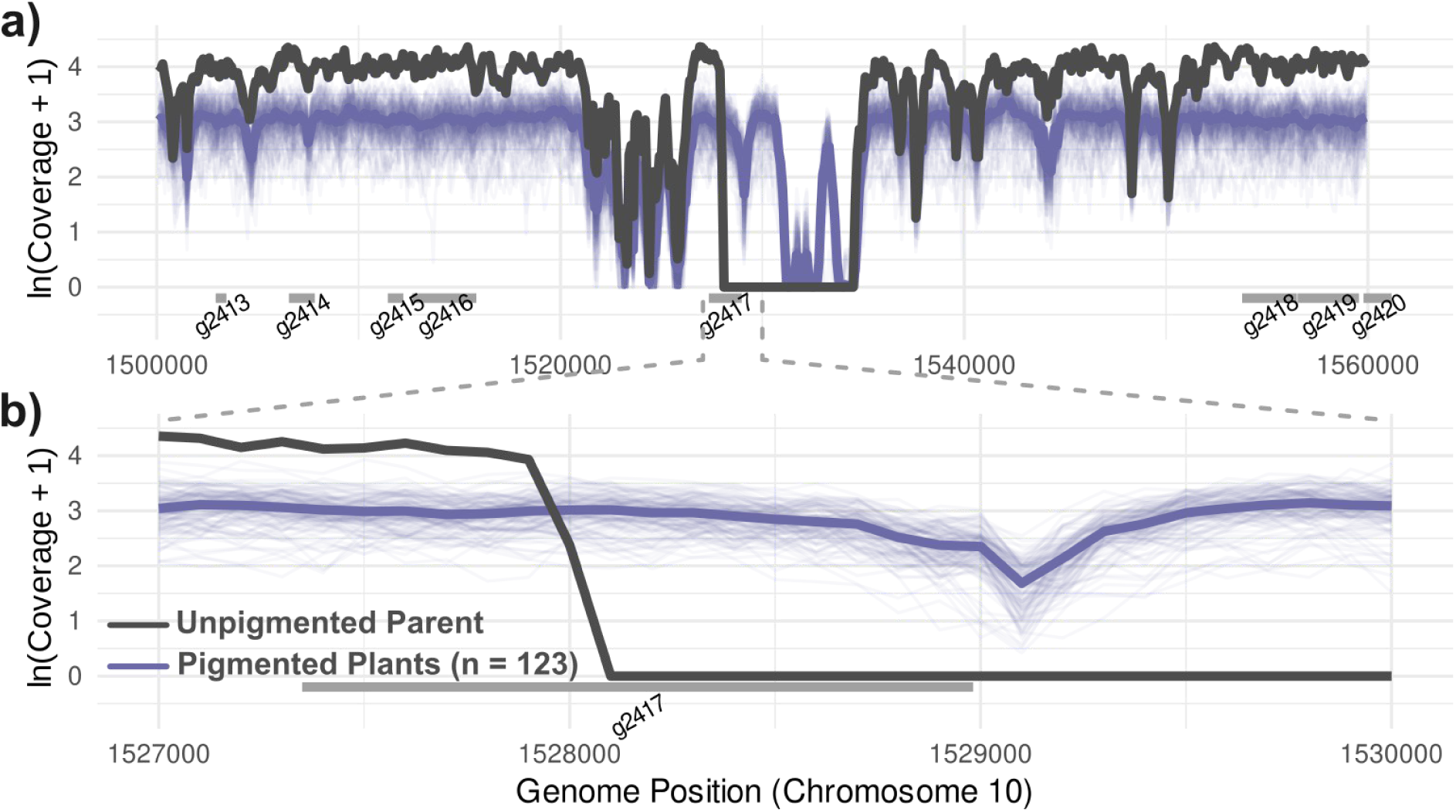
*IhUF3GT1* deletion is unique to unpigmented plants. a) The locus containing the first half of *g2417* (encoded on the minus strand) is absent in unpigmented plants. Of 123 pigmented lines, all pigmented plants (123/123) contain the coding sequence of *g2417*, including the start codon. Log-transformed read counts (i.e., coverage) of 100 bp windows in each genotype. Solid grey line is coverage in the unpigmented genotype. Solid purple line is the average coverage in all 123 pigmented plants. Faint purple lines are coverage in each pigmented plant. Coverage data for pigmented lineages is from (Peake *et al*., 2026). b) Zoomed view of coverage in *g2417* specifically.

*IhUF3GT1* (*g2417*) also perfectly co-segregates with the pigmented phenotype. We determined if unpigmented plants in an F_3_ population lacked *IhUF3GT1* using endpoint PCR and electrophoresis. We found that all pigmented plants in our co-segregation analysis contained the *IhUF3GT1* sequence (36/36; Figure 5). No unpigmented plants contained the *IhUF3GT1* sequence (0/17; Figure 5). Taken together, our results demonstrate that the deletion of the *IhUF3GT1* locus underlies the absence of pigments in plants with white flowers.

**Figure 5.**
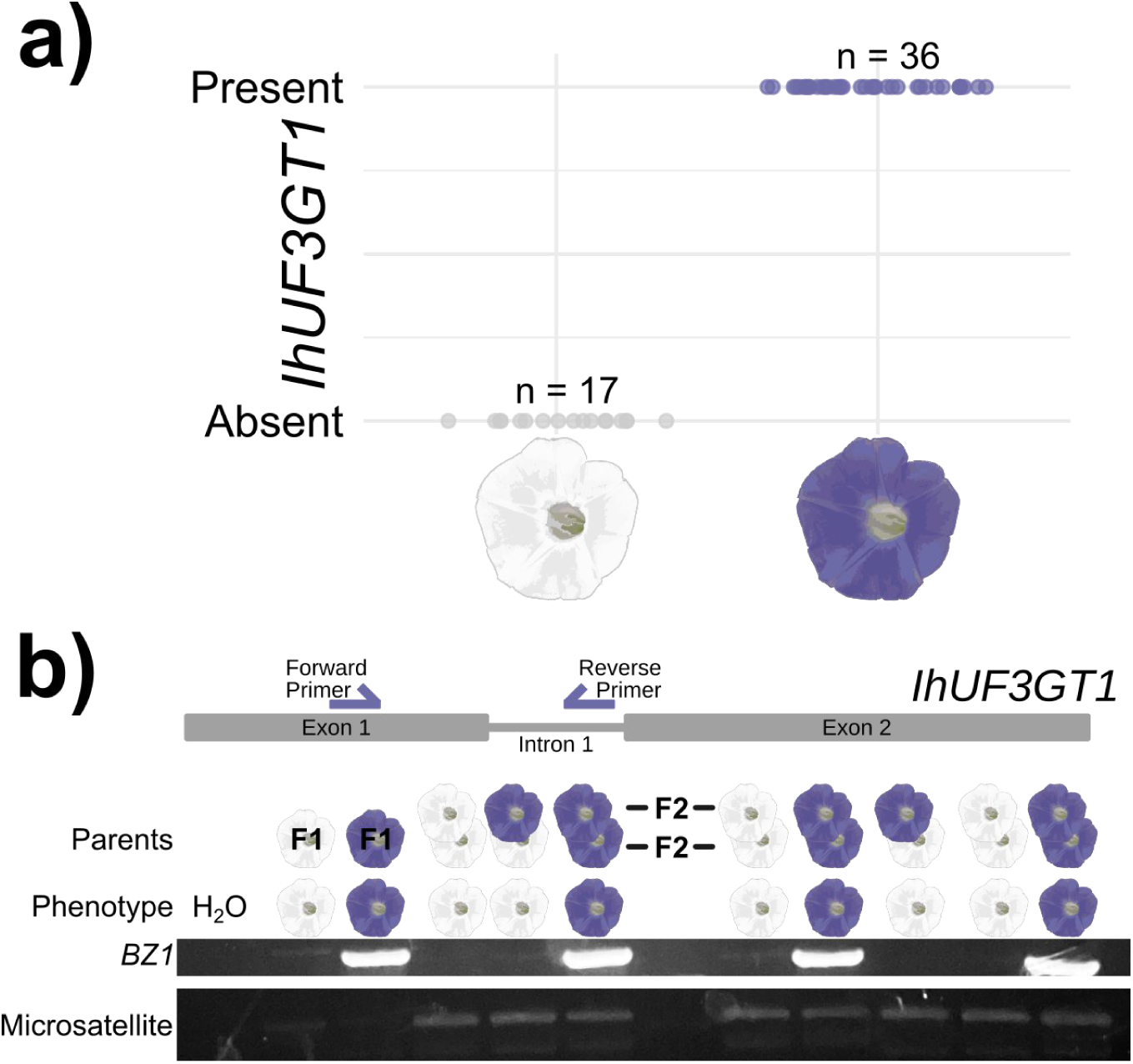
Deletion of *IhUF3GT1* perfectly co-segregates with the unpigmented phenotype. a) Summarized dot plot of endpoint PCR detecting the *IhUF3GT1* deletion in F_3_ plants of unpigmented and pigmented plants. Grey points represent unpigmented plants. Purple points represent pigmented plants. Detection of *IhUF3GT1* is based on the visibility of a band on an agarose gel. *IH00534 (Campitelli, and Stinchcombe, 2014)* is a universal microsatellite marker for *I. hederacea* and serves as reaction positives for all samples. Sterile water served as a reaction negative. A full sibling cross of the unpigmented parent served as the biological negative. A full sibling cross of the pigmented parent served as the biological positive. b) Example gel used to determine presence/absence of *IhUF3GT1*. Presence/absence of *IhUF3GT1* is related to the phenotype in the individual with schematic of primer positions on *IhUF3GT1* genomic sequence (exact positions in Table 1).

## DISCUSSION

We characterize a phenotypically and genetically rare phenotype in Ivy-leaf morning glory (*Ipomoea hederacea*), a complete lack of anthocyanins due to a mutation in which flowers are completely white and no purple pigmentation is found elsewhere on the plant. By combining test crosses, transcriptomics, genome coverage, and co-segregation analyses, we identify the genetic basis of this new, unusual phenotype. Loss of pigmentation in *I. hederacea* is determined by a heritable, recessive deletion of *anthocyanidin 3-O-glucosyltransferase* (*IhUF3GT1*). We propose that these unpigmented *I. hederacea* plants are unable to convert cyanidin into stable blue/purple anthocyanin pigments because they lack *IhUF3GT1*. Below, we 1) discuss how the *IhUF3GT1* mutant is a unique model to test the fitness effects of anthocyanins specifically (i.e., in the absence of pleiotropic effects) and 2) hypothesize that anthocyanin loss is sufficiently deleterious that selection likely favors pigmented plants in natural populations, leading to the maintenance of pigmented phenotype in this species.

Anthocyanin biosynthesis genes are highly conserved across plants (Rausher *et al*., 1999; Saigo *et al*., 2020; Buitrago *et al*., 2025), so much so that anthocyanin biosynthesis genes are an excellent model for comparative genomics among highly diverse plant lineages (Piatkowski *et al*., 2020; Buitrago *et al*., 2025). There has been a wide-spread prediction that mutations in genes early in the anthocyanin pathway may be highly pleiotropic because virtually every step preceding anthocyanin biosynthesis creates precursors for other branches of flavonoid biosynthesis. Thus, mutations in most anthocyanin biosynthesis genes may extend beyond just alterations in anthocyanins themselves but also impact flavonols (Daryanavard *et al*., 2023), isoflavonoids (Wang *et al*., 2025), proanthocyanidins (Furukawa *et al*., 2007), etc. One consequence is that any loss-of-function mutations upstream of cyanidin/pelargonidin synthesis will affect both anthocyanins and proanthocyanidins (and potentially other secondary metabolites), making it difficult to disentangle their fitness effects. However, glycosylation of anthocyanidins by *IhUF3GT1* is the final step in anthocyanin biosynthesis (Rausher, 2006). The mutant we have characterized is thus a potentially useful tool for studying the fitness consequences of anthocyanin loss specifically, because we do not expect the deletion of *IhUF3GT1* to have pleiotropic consequences on traits derived from other branches of the pathway. *Ipomoea hederacea* is a tractable species for field experiments (Campitelli and Stinchcombe, 2013a; Henry and Stinchcombe, 2023b, 2025; Boyle *et al*., 2024; Stinchcombe 2002), suggesting that the F_2_ and F_3_ populations we developed could be used for testing the fitness consequences of anthocyanin loss, in an experimental population with much of the genetic background randomized by recombination.

More broadly, loss of pigmentation in *I. hederacea* is far rarer than in its close relative *I. purpurea* (Ennos and Clegg, 1983; Coberly, 2003; Coberly and Rausher, 2008). The *IhUF3GT1* mutant we have discovered and identified is the first *I. hederacea* sample we have observed with unpigmented flowers (in >25 years of field work on the species and out of ∼750 naturally collected lines in our lab; Stinchcombe, personal observations). Other researchers also report unpigmented phenotypes to be rare in this species: Dr. Regina Baucom reports encountering a single unpigmented individual across four years of field surveys of ∼ 50 populations each throughout the Southeastern and Midwestern portion of the range, comprising thousands of individuals in field settings (RS Baucom, personal communication) In addition to the phenotypic rarity of complete anthocyanin loss in *I. hederacea*, the genetic basis of our unpigmented line is also genetically rare because no other sequenced line to date (0/123) has a *IhUF3GT1* deletion. If, as described above, mutations at the end of the anthocyanin pathway are predicted to have weaker deleterious pleiotropic effects on other traits, why is pigmentation loss so rare in *I. hederacea*, especially compared to the closely related *I. purpurea*? We suggest two possible explanations for the differences. First, the *w* and *a* alleles in the mixed-mating *I. purpurea* appear to be protected from loss due to a transmission advantage associated with increased selfing when homozygous (*w*, *Brown and Clegg, 1984; Rausher et al., 1993*; *a*, *Fehr and Rausher, 2004)*.

Because selfing rates are much higher in general in *I. hederacea* (often >90%), it seems unlikely that they would produce fitness differences between blue and white-flowered *I. hederacea,* and contribute to the maintenance of the *IhUF3GT1* deletion. A second explanation of the rarity of *IhUF3GT1* deletions is simply that selection is acting against anthocyanin loss and thus the frequency of the *IhUF3GT1* deletion is governed by mutation-selection balance. The overall rarity of whole plant anthocyanin loss (Del Valle *et al*., 2019) and loss-of-function mutations to the enzymatic/structural genes of the anthocyanin pathway (Wu *et al*., 2013; Coburn *et al*., 2015), are consistent with this hypothesis. However, in contrast to the *a* locus in *I. purpurea*, and the *DFR* loss-of-function mutations in *Mimulus lewisii* and *Iochroma calycinum*, we hypothesize that the rarity of unpigmented plants is due to selection against anthocyanin loss *per se*, rather than its pleiotropic consequences.

In short, we discover a monogenic *IhUF3GT1* mutant in *I. hederacea*. This *IhUF3GT1* mutant is an excellent tool to disentangle the ecological consequences of anthocyanin loss from loss of “linked” traits and to study how selection acts on a single locus affecting an ecologically important trait. More broadly, the rarity of both anthocyanin loss and loss-of-function in *IhUF3GT1* is highly indicative of the strong selective force of anthocyanins in this species. Consequently, the *I. hederacea IhUF3GT1* loss-of-function mutant can serve as a critical model system to understand how anthocyanins specifically may shape the fitness and selective forces underlying economically significant crops (e.g., sweet potato), crop-destroying weeds (e.g., *I. hederacea*), and ornamental plants (e.g., *I. purpurea*) in the *Ipomoea* genus. This *IhUF3GT1* loss-of-function mutant can allow for identifying and quantifying the mechanism by which anthocyanins determine fitness in *I. hederacea* and perhaps other *Ipomoea* species such as through UV protection, drought resistance, and defense against herbivores.

## Acknowledgments

We thank Katie Maunder and the rest of the Stinchcombe lab for comments and discussion on the manuscript, and Thomas Gludovacz and Alice DesRoches for horticultural support, and 3 anonymous reviewers for comments on the manuscript. Funding from NSF (DJH: DBI-2305481), an NSERC Discovery Grant (JRS: RGPIN-2022-04366), NSERC Graduate Fellowships (ALP), and the EEB Department (ML, MRH) supported our work. We thank Mark Rausher for suggesting the RNA-Seq experiment, and Stacey Smith and Sylvie Martin-Eberhardt for suggestions. We thank Regina Baucom for sharing how often she had encountered a similar phenotype in field surveys. Our research was enabled by support from Compute Ontario and the Digital Research Alliance of Canada.

## Author Contribution Statement

Conceptualization and Supervision: JRS. Laboratory Experiments and Plant Growth: EG, MCL, MRH, ALP. Computational Analyses: DJH, ALP. Funding Acquisition: DJH, JRS, ALP, MCL, MRH. Writing: DJH, EG, JRS. Visualization: DJH.

## Data Archiving

Gene expression and genomic data from the unpigmented parent are deposited in NCBI (BioProject: PRJNA1355176, SRA: SRP638260). Our code and computational environments will be posted on GitHub following publication. Packages used to conduct analyses in R (v4.5.1; R Core Team, 2025) are also listed in Supplementary File 1.

## Research Ethics Statement

This work did not involve humans, animals, their data, or their biological material.

---

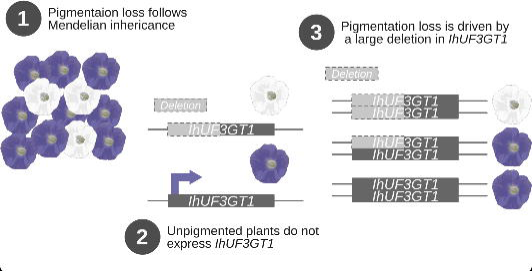

